# Population genomic analysis reveals cryptic population structure in the commercially important Lake Malawi cichlid *Copadichromis mloto*

**DOI:** 10.1101/2024.04.26.591058

**Authors:** Wilson Sawasawa, Alexander Hooft van Huysduynen, Sophie Gresham, Harold Sungani, Bosco Rusuwa, Gudrun De Boeck, George F. Turner, Maxon Ngochera, Hannes Svardal

## Abstract

Fish is an important source of animal protein for many people living around Lake Malawi. The evaluation of population structure and genetic diversity can yield useful information for management and conservation of fish species but is complicated in Lake Malawi by the close genetic relatedness of species of the recent cichlid adaptive radiation. In this study, we analysed whole-genome sequencing data of “true utaka”, a group of cichlids previously common in fisheries, but that has faced strong decline due to overfishing. Our analysis of 223 individuals collected from fishermen’s catches along the western shoreline of Lake Malawi confirmed that *Copadichromis mloto* (*C. sp*. “virginalis kajose”) is the true utaka most targeted by fisheries. Genetic principal component analysis, phylogenetic inference, and admixture analysis revealed complex patterns of population structure. The presence of at least three geographically widespread genetic clades that have remained separate despite gene flow and partial sympatry hints at the presence of currently undescribed, cryptic species diversity in *C. mloto*. This result leads us to suggest that, despite the lack of obvious habitat barriers, benthic and pelagic species of Malawi cichlids might harbour unidentified species diversity and calls for further genetic and taxonomic research to define appropriate conservation units.

## Introduction

Cichlid fishes of the Great Lakes of East Africa, particularly Lakes Victoria, Malawi, and Tanganyika are significant study systems in evolutionary biology because of their rapid adaptive radiation (Anseeuw et al. 2008; Brawand et al. 2015; Rick et al. 2022). At the same time, Malawi cichlids are a source of food for millions and provide a livelihood for thousands of rural and urban Malawians who participate in harvest and postharvest activities including fishing, fish processing, transportation to the markets, and marketing (Weyl et al. 2004). Stocks of Malawi cichlids have been dramatically declining over the past 40 years, probably as a consequence of this exploitation (Van Zwieten et al. 2011; Ngochera et al. 2018; Chavula et al. 2023), calling for sustainable fisheries management strategies. However, monitoring of exploited populations has been severely constrained by difficulties in species identification, a particular challenge for the Lake Malawi cichlid radiation with its ∼1000 closely related and taxonomically poorly resolved species (Svardal et al. 2020). Cryptic species diversity can mask changes in community composition, interactions between different fishery sectors and gears and render studies of population structure unreliable, as apparent genetic differentiation may instead represent non-random sample mixes of different species collected at different locations.

Previous studies on the genetic structure of Malawi cichlids suggest that geographic distance and habitat barriers contributed to population structure in rock-dwelling species such as the species-rich clade of Mbuna (Van Oppen et al. 1997; Danley et al. 2000) or *Protomelas* (Pereyra et al. 2004), but found no or only weak population structure within the more continuously distributed species living over sandy bottom or in the open water (Shaw et al. 2000; Taylor & Verheyen, 2001; Anseeuw et al. 2011), which constitute the main target of fisheries. However, traditional genetic markers as used in these studies (mtDNA and microsatellites) may lack power to identify cryptic species (Malinsky et al. 2018), which can lead to misleading population structure estimates. Conversely, recent whole genome studies on Malawi cichlids have proven powerful in resolving species relationships (Malinsky et al. 2018; Masonick et al. 2022; Scherz et al. 2022), but were limited to one or few samples of each species and thus not able to resolve geographic structure within species. As such, we lack a detailed understanding of the patterns of population structure of Malawi cichlid species.

The semi-pelagic utaka (*Copadichromis sp*. (Turner, 1996) have traditionally been among the economically most important fish species in Lake Malawi and Lake Malombe, a highly productive and heavily fished lake at the outflow of Lake Malawi (Tweddle et al. 1995; Weyl et al. 2004; Kanyerere et al. 2018). Among the most common Utaka in fishery catches are a group known as the ‘pure utaka’ which lack dark spots on their flanks, described as *C. mloto* (slender-bodied) and *C. virginalis* (deeper bodied), but long recognised as posing considerable taxonomic and identification issues (Iles, 1960; Eccels & Trewavas, 1989). The ‘pure utaka’ most common in the fisheries of both lakes, previously often referred to as *C*. sp. ‘virginalis kajose’, has recently been re-identified as *C. mloto* (Turner et al. 2022). This species seems to be ecologically differentiated from *C. virginalis*, (previously referred to as *C*. ‘virginalis kaduna’; *C. ilesi*; *C*. sp. ‘firecrest mloto’) by its habit of breeding over the open sandy or muddy substrate, rather than on rocky coast (Turner et al. 2022). The stocks of *C. mloto* have dramatically declined in Lakes Malawi and Malombe over the past decades, likely as a result of overfishing (Banda et al. 2001; Weyl et al. 2004; Anseeuw et al. 2011; Chavula et al. 2023).

In this study, we analysed whole genome sequences of 203 specimens of *C. mloto* collected with a research trawler and from fisheries catches along the Malawian waters of Lake Malawi and from Lake Malombe, covering a latitudinal extent of more than 480 kilometres. We confirm the monophyly of *C. mloto* with respect to *C. virginalis*. Analysis of population structure revealed the presence of three genetically distinct but geographically overlapping clades within *C. mloto*, hinting at the presence of hitherto cryptic taxonomic diversity, yielding important information for the conservation and management efforts of these populations. Gene flow analysis confirmed historic genetic exchange both between *C. virginalis* and *C. mloto* and among the different *C. mloto* clades, the latter showing correlation with geographic proximity. Furthermore, we found similar levels of genetic diversity across *C. mloto* populations and a strong excess of rare genetic variants consistent with historic population growth but could not detect any discernible effect of fishing pressure on population genetic statistics. Finally, we discuss tentative evidence for differences in male breeding colouration between the observed genetic clades.

## Materials and Methods

### Sample collection

Specimens of *Copadichromis mloto* (n = 202) *and C. virginalis* (n = 21) were collected along the western shoreline of Lake Malawi, its southeast and southwest arms, and Lake Malombe (Supplementary Table 1, Fig. 1) during field expeditions in 2002 (Anseeuw et al. 2011), 2016, and 2017 (Supplementary Table 1). The samples were caught by local fishermen using a variety of methods, including beach seine nets, open water seines (Chirimila nets in Lake Malawi), purse seines (Nkacha nets in Lake Malombe) and trawl ships (both commercial and the Ndunduma research trawler of the Malawi Department of Fisheries) (Supplementary Table I). The specimens’ right pectoral fins were preserved in pure ethanol and stored in a freezer. A subset of whole fish specimens was fixed in 10% formalin and preserved in ethanol; specimens are being kept at the Royal Museum for Central Africa in Tervuren, Belgium (2002 samples) or the Zoology Museum Cambridge (2016-2017 samples). However, we note that for the specimens stored in the Royal Museum for Central Africa no 1:1 match to fin clips is available.

**Figure 1.**
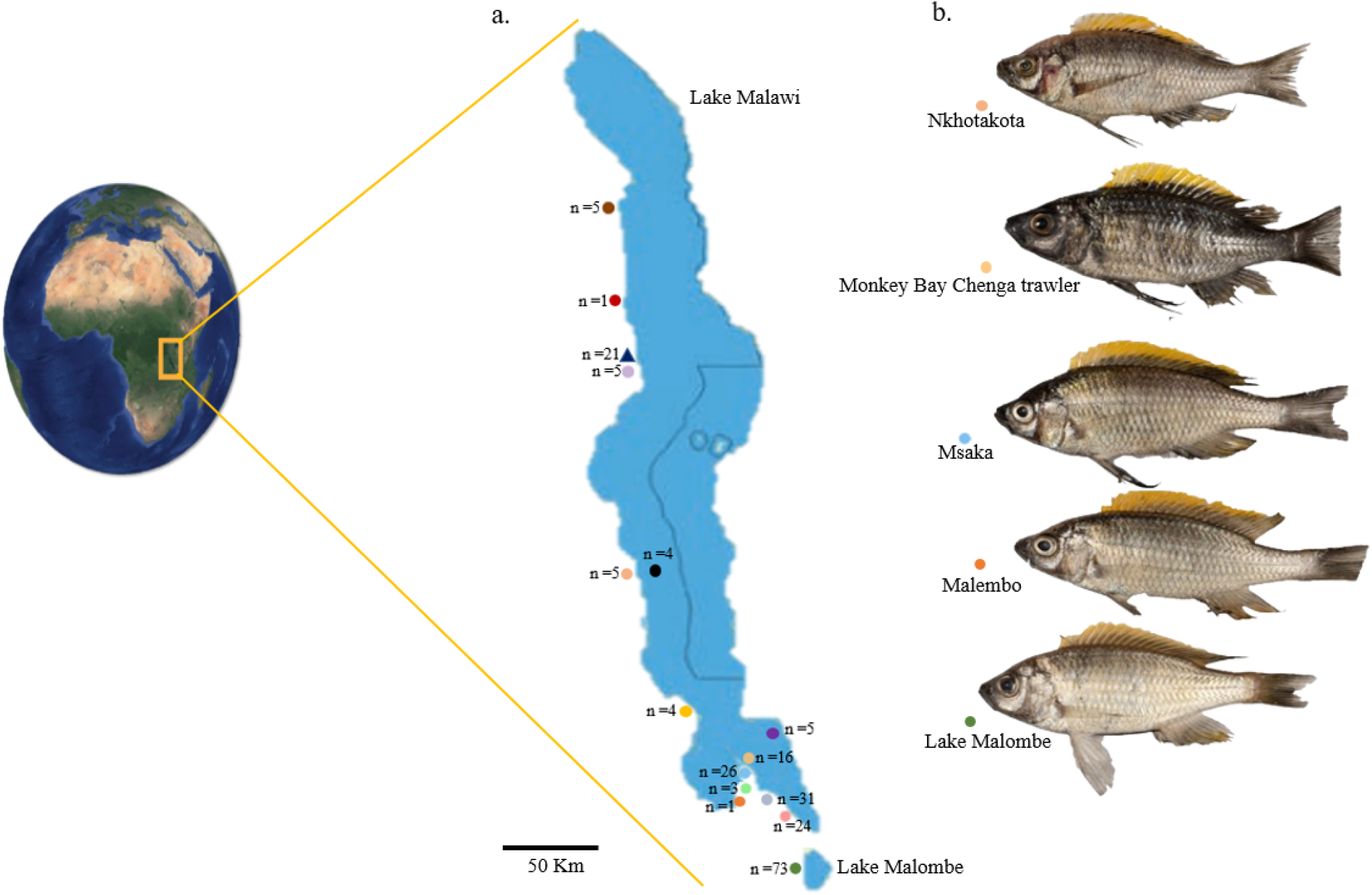
Sampling information. (a) Location of Lake Malawi on the African continent (left) and zoom in of Lakes Malawi and Malombe (right), showing the number of samples and sampling locations of *Copadichromis mloto* (circles) and *C. virginalis* (triangle). Location names are given in Fig. 2A and Supplementary Table 1. (b) Representative photos of males in breeding colouration from the locations Nkhotakota, Monkey Bay Chenga trawler, Msaka, Malembo and Lake Malombe (top to bottom).

### Sequencing, variant calling and filtering

DNA was extracted and libraries were prepared at the Wellcome Sanger Institute using standard protocols. Whole-genome sequencing was performed on an Illumina HiSeq platform to individual coverages ranging from 7.2-fold and 42.1-fold. Resulting short read sequences were aligned to the *Astotilapia calliptera* reference genome fAstCal1.2 ((GCA_900246225.3 https://www.ncbi.nlm.nih.gov/assembly/GCF_900246225.1) using BWA-MEM (BWA version 0.7.17). Furthermore, bcftools (version 1.14) was used to detect variant sites (Danecek et al. 2021). Sites were masked if the overall mapping quality was less than 50 or if more than 10% of the mapped reads had a mapping quality of zero, diagnostic of non-unique mapping. In addition, sites for which mapping quality was significantly different between the forward and reverse strand (P<0.001) were also masked, as well as sites for which the sum of overall depth for all samples was unusually high (>97.5 percentile) or low (<2.5 percentile). Heterozygous genotypes for which a binomial test showed a significantly biased read depth between the reference and alternative alleles (PHRED score >20) were set to missing, representing between 0.02% and 0.15% of heterozygous sites per individual. Additionally, sites with excess heterozygosity (dataset-wide inbreeding coefficient < 0.2) and sites with >20% missing genotypes were excluded from analysis (Supplementary Figs VI, VII and VIII). Only biallelic single nucleotide polymorphisms (SNPs) were retained for further investigation, while recently discovered inversion regions on five chromosomes (total 104 megabase pairs [Mbp]) were excluded from further analysis (Supplementary Table II, Blumer et al. in preparation). Furthermore, we excluded ten samples because they showed abnormally high heterozygosities reminiscent of cross contamination, or were inferred to be close relatives.

### Population genetic structure

For this analysis, we only considered SNP variants with a minor-allele count of at least 3 and less than 25% missing genotypes filtered for linkage-disequilibrium in PLINK (version 1.9) with r^2^ > 0.2 in sliding windows of 50 Single Nucleotide Polymorphism (SNPs), with 10 SNP overlap. We used PLINK to run a principal component (PC) analysis calculating eigenvectors and eigenvalues separately for the whole dataset, including *C. virginalis* and *C. mloto*, and for *C. mloto* only (Purcell et al. 2007). R (version 4.2.2) (R Core Team, 2021) was used to plot the eigenvectors.

### Population admixture analysis

We used the LD pruned sites mentioned above to create binary PLINK (.bed) files. The .bed files were used to perform admixture analysis using Bayesian clustering in ADMIXTURE v1.3.0 (Alexander et al. 2009) with values of *K* ranging from 1 to 5. We selected the cluster K value with the lowest corresponding CV error. Python (version 3) (Van Rossum & Drake, 2009) was used to plot the results of the admixture analysis. The admixture analysis results were plotted and visualised using the Pong workflow analysis (https://github.com/ramachandran-lab/pong).

### Neighbour-joining tree construction

To construct a neighbour-joining (NJ) tree of genetic distances, we calculated total pairwise genetic differences on the genome-wide SNP loci of the samples using a custom script based on Scikit allel and the Biophyton 1.79 phyllo package for tree construction (https://github.com/feilchenfeldt/pypopgen3). The tree was rooted using an ancestral sequence inferred by whole genome alignment of the reference genome and outgroup species (Camacho et al. in prep.).

### Population genetic statistics

Genetic diversity statistics were calculated for populations of *C. mloto* and for the single population of *C. virginalis* from Nkhata Bay. The inbreeding coefficient (F) per individual, and nucleotide diversity per site and Tajima’s D in windows of 100kbp were calculated using vcftools (version 0.1.16) (Danecek et al. 2021). Furthermore, the allele frequency spectra were calculated for the population of each *C. mloto* clade with the largest sample size using a custom script.

### Gene flow test in *Copadichromis* species

To investigate potential gene flow between *C. virginalis and C. mloto* and between different *C. mloto* populations, we computed four population tests such as Patterson’s *D* and the *f*_*4*_ (admixture) ratio (also known as the ABBA-BABA statistics) as implemented in D-suite (Malinsky et al. 2021), setting the above-mentioned ancestral sequence as outgroup. We defined as populations all the samples of a given sampling location assigned to one of the identified genetic clades (*mloto A/B/C*, putatively admixed, *C. virginalis*). For analysis, we selected all *f*_*4*_ ratio tests comparing populations from specific clades, e.g., P1 = population from clade *mloto A*; P2 = population from clade *mloto B*; P3 = *C. virginalis*, etc. Finally, to assess geographic signal in *f*_*4*_ ratio tests we obtained sampling coordinates either from available field records or, where not available, by locating reported sampling sites on online maps, and then calculated geographic distances using the function distance.geodesic implemented in the Python package geopy (version 2.3.0).

## Results

### Variant detection

Variant calling yielded approximately 41 million single nucleotide polymorphisms (SNPs) across 21 *C. virginalis* and 203 *C. mloto* specimens of which ∼31 million (74.7%) passed all variant filtering criteria. The accessible genome size excluding recently discovered inversion regions (see methods) was 545 Mbp corresponding to a SNP density of 75 SNPs per kbp.

### Population genetic structure provides evidence of three geographically widespread *C. mloto* clades

Performing principal component (PC) analysis and constructing a neighbour-joining (NJ) tree of pairwise differences of all samples, we confirmed that all *C. mloto* formed a monophyletic clade with respect to the samples identified as *C. virginalis* (Supplementary Fig. V). The first PC clearly separated *C. mloto* samples from all sampling locations from the single *C. virginalis* population from Nkhata Bay (10 % variance explained). Since our collections from fisheries catches did not initially distinguish between the two species, this result suggests that *C. mloto* is the dominant pure utaka species in commercial and artisanal fisheries of Lakes Malawi and Malombe. For further analyses, we focused on the 203 specimens belonging to *C. mloto*.

Structure/admixture analysis of *C. mloto* samples revealed an interesting pattern in that identified ancestries did not perfectly reflect geographic proximity. Although most individuals of the same sampling location were attributed to the same cluster, the different clusters spanned a wide geographic range with individuals from non-adjacent sampling locations being attributed to the same cluster, while geographically intermediate populations were attributed to different clusters (Fig. 2A This trend was consistent for different choices of ancestral populations (K) (Supplementary Fig. IV). For several populations, some individuals were inferred to draw some or all of their ancestry from a cluster different from the majority of individuals, consistent with the presence of distinct local populations and, in the case of partial ancestry, gene flow, which will be further investigated below.

**Figure 2.**
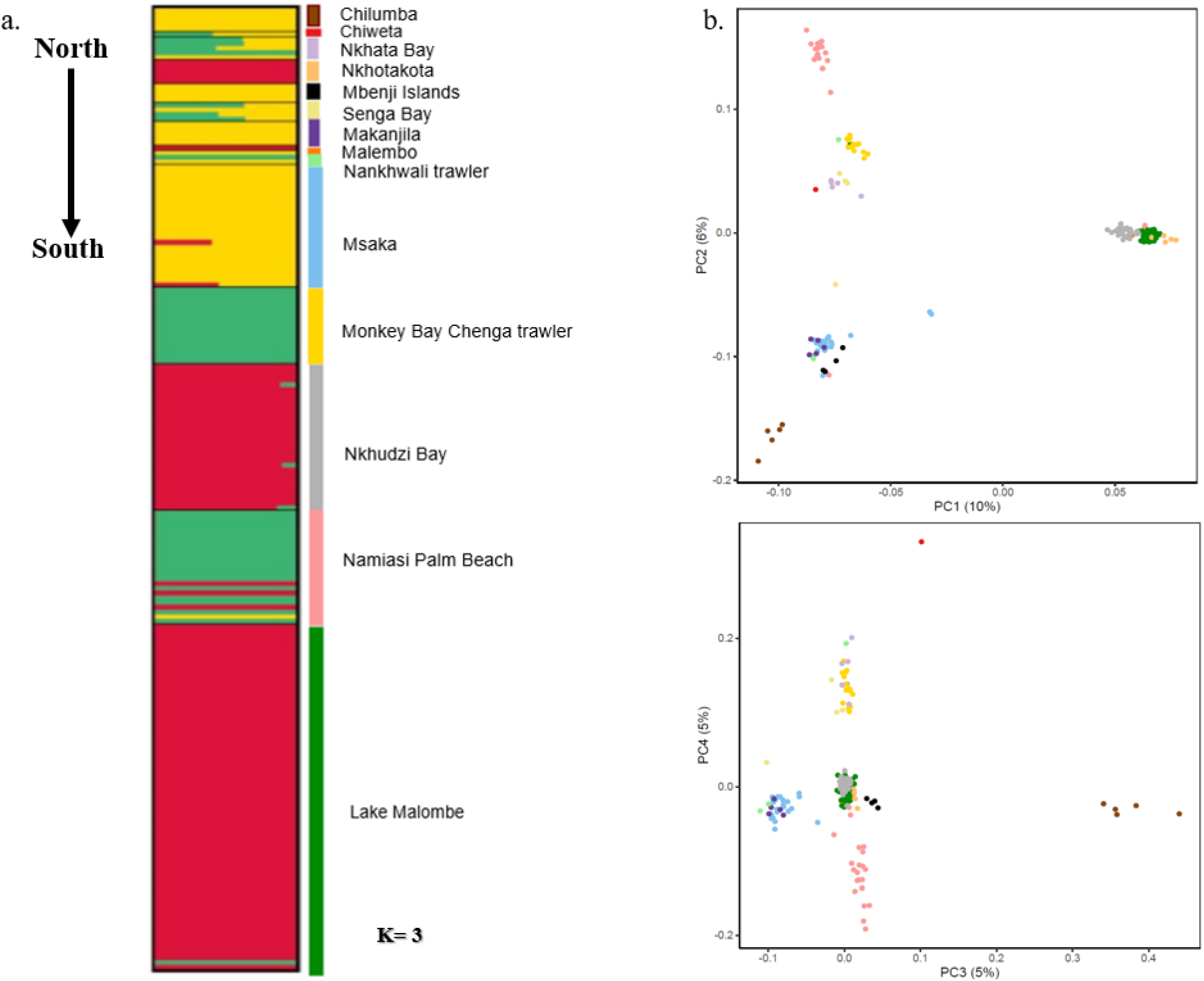
Relationships among the *Copadichromis mloto* populations visualised by population Admixture and Principal Component (PC) analysis. (a) Population structure patterns of *C. mloto* as inferred by Admixture (Alexander et al. 2009) assuming K = 3 ancestral populations; for the other K values see Supplementary Fig. IV. (b) Principal Component (PC) analysis of *C. mloto* populations.

A PC analysis of *C. mloto* individuals broadly supported the results of the admixture analysis in that it identified genetic clustering inconsistent with a simple isolation by distance pattern (Fig 2B). Instead, the first PC separated a clade consisting of most Lake Malombe individuals, all individuals from Nkhudzi Bay on the western shoreline of the southeast arm of Lake Malawi, all individuals from Nkhotakota on the central western shore of Lake Malawi, some individuals from Namiasi Palm Beach at the southernmost tip of the south east arm of Lake Malawi as well as a single individual from Malembo in the south west arm of Lake Malawi. These individuals formed a monophyletic sister clade to all other individuals in an NJ tree of *C. mloto* specimens (Fig. 3). In the following, we refer to this genetic cluster as *mloto A* clade.

**Figure 3.**
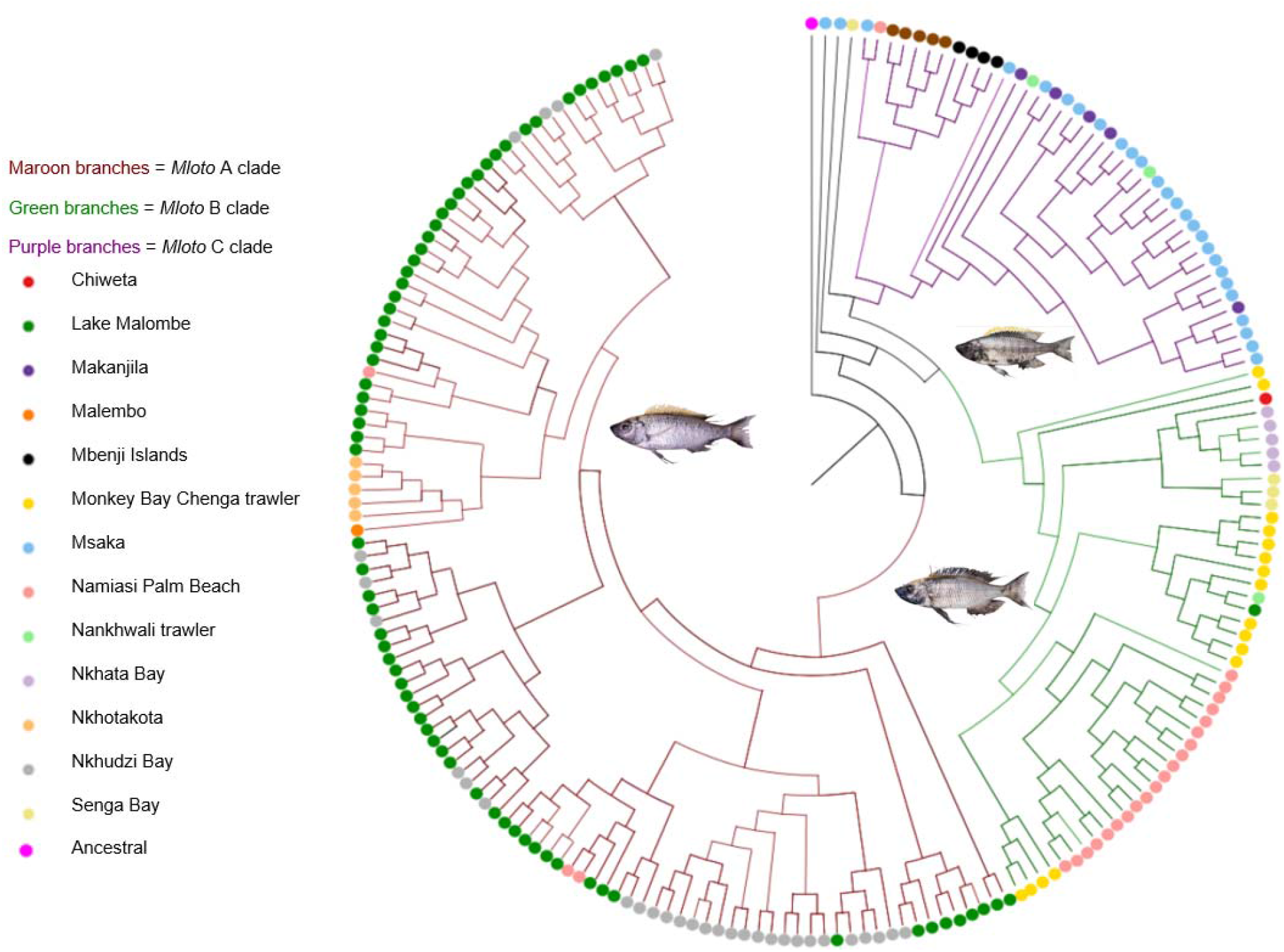
Neighbour-joining tree constructed from pairwise distances showing the three genetic clades of *C. mloto*. The branches corresponding to the three genetic clades are coloured in red (*mloto A*), green (*mloto B*), and purple (*mloto C*), respectively. Black branches correspond to the ancestral sequence used for rooting and to putatively admixed individuals. Relative branch length is fixed for better visibility and thus not proportional to genetic distance (See Supplementary Fig. XII for a tree with distances). Representative photos of breeding males from each genetic clade are given (Fig. 6).

The second PC separated samples more gradually, with individuals from the northern most population at Chilumba and most individuals from the Namiasi Palm Beach population at the southern tip of Lake Malawi at the two extremes and the *mloto A* clade that separated along PC1 at values close to zero. However, the clustering of other populations did not reflect a clear geographic trend: individuals with a wide range of geographic origins clustered next to the Namiasi Palm Beach cluster, including samples from Chiweta, Nkhata Bay, and Senga Bay along the western shore of Lake Malawi as well as individuals from the southeast and southwest arm populations. Conversely, individuals from other south west arm populations, as well as from Makanjila on the south eastern coast of Lake Malawi, and a single sample from Namiasi Palm Beach, clustered in proximity to the northern most samples from Chilumba. In the NJ tree, the described spread along PC2 corresponds to two reciprocally monophyletic clades, which we refer to as clades *mloto B* and *mloto C*, in the following, for the clades including samples from Chilumba (negative PC2) and Nkhata Bay (positive PC2), respectively. An exception to this are two samples from Msaka on the eastern shores of the southwest arm that clustered basally to both *mloto B* and *mloto C* and intermediate along PC2. This is consistent with admixed ancestry between the clades, as also inferred by Admixture (Fig. 2A), and further investigated below.

### Differential genetic exchange between *Copadichromis virginalis and C. mloto* populations

To test whether the evolutionary separation between *C. virginalis* and *C. mloto* corresponded to a clean bifurcation or whether *C. virginalis* and the identified subclades and populations of *C. mloto* continued to exchange genetic material, we calculated the f4 admixture ratio – a measure of excess allele sharing of a clade P2 with a clade P3, relative to P2’s sister clade P1. Specifically, we estimated a potential contribution of *C. virginalis* to a given *C. mloto* population compared to another *C. mloto* population, where we defined populations as all samples of a specific sampling location that fall into the same *C. mloto* clade as identified above. The test revealed strong (5-25%) and highly significant excess allele sharing of all *mloto C* populations with *C. virginalis* compared to all *mloto A* populations and most *mloto B* populations (Fig. 4a). Furthermore, the *mloto C* populations from Chilumba, Mbenji Islands, and Namiasi Palm Beach showed 9-10% excess allele sharing with *C. virginalis* compared to other *mloto C* populations. Finally, *mloto B* populations showed 4-10% excess allele sharing with *C. virginalis* compared to *mloto A* populations. Overall, these results suggest differential exchange of genetic material between *C. virginalis* and different *C. mloto* populations, with the highest levels of *C. virginalis* contributions in some *mloto C* populations and the lowest in *mloto A*.

**Figure 4.**
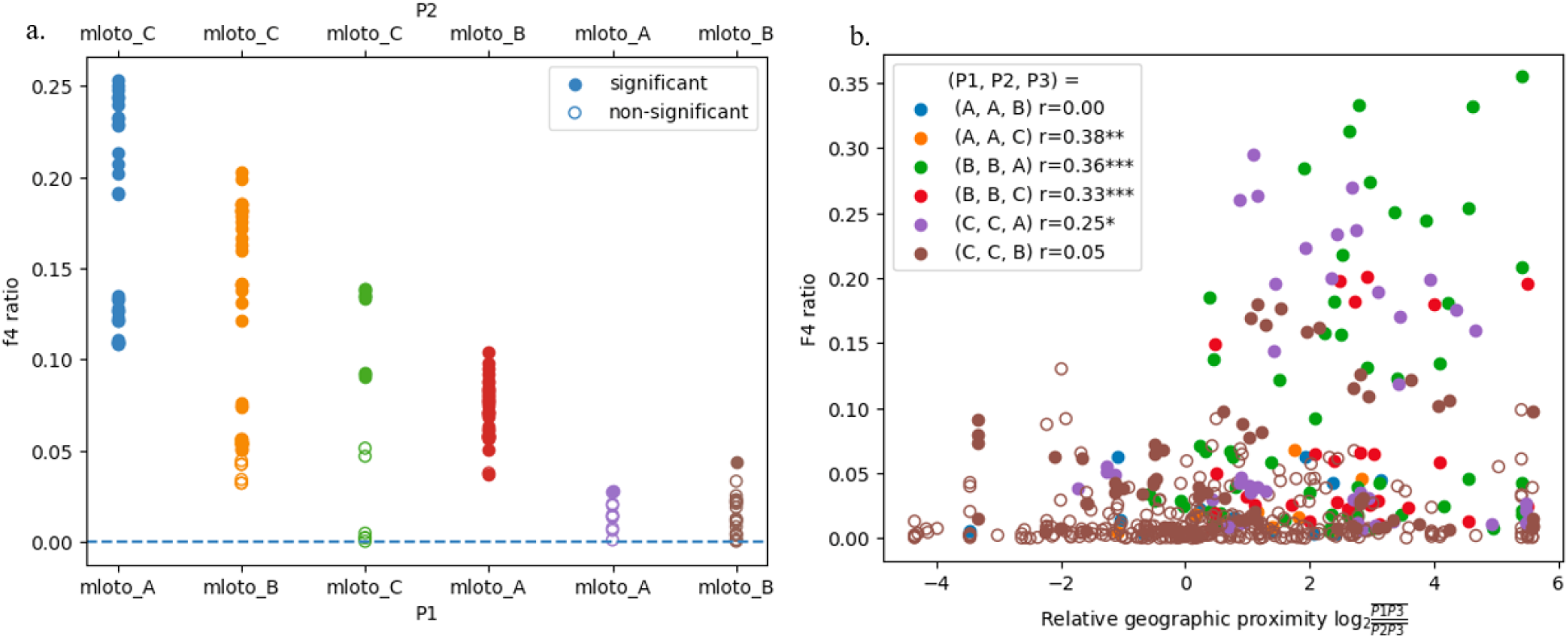
Dtrios for gene flow test in *Copadichromis mloto* and *C. virginalis* populations. (a) f4 admixture ratio tests of the form f4 (P1, P2, P3, Outgroup) in which P1 and P2 are *C. mloto* populations and P3 *C. virginalis*, segregated by *C. mloto* clade adherence of P1 and P2. (b) f4 admixture ratios plotted against relative geographic proximity among the *Mloto* clades. Points are coloured by *C. mloto* clade adherence of P1, P2, P3 populations in tests f4 (P1, P2, P3, Outgroup) (See Methods).

### Evidence for local genetic exchange between *C. mloto* clades

Next, we wanted to test whether the three *C. mloto* clades are fully reproductively isolated entities or whether gene flow has occurred between the clades after their initial split. In the former case of clean splits between the clades without subsequent gene flow, we would expect that individuals from two given clades are equally closely related to each other irrespective of their geographic origin, while in the latter case of cross-clade genetic exchange, we would expect cross-clade excess allele sharing of the populations involved, meaning that different populations of one clade show variation in their genetic distance to an outgroup population. We first checked for signals of excess allele sharing of the two samples from Msaka which clustered basally to the *mloto B* and *C* clades and found that these samples showed highly elevated f4 admixture ratios of up to 42% with *mloto A* populations relative to most *mloto B* and *C* populations, while also generally closer to *mloto B* than to *mloto C* populations. This provides further evidence for the admixed status of these two samples with ancestries related to *mloto A* and *mloto B* clades as already suggested by the Admixture analysis.

Overall, we found strong evidence for genetic exchange between the different *C. mloto* clades with 49% of f4 ratio tests significant above multiple testing (Bonferroni FWER < 0.05). If this pattern were due to relatively recent gene flow, we would expect a trend in which individuals from different clades are relatively more closely related to each other if they originate from nearby sampling locations. To investigate this, we tested whether excess allele sharing between populations of different clades depended on their relative geographic distance. We found that most clade comparisons showed a significantly positive correlation between relative geographic proximity and signatures of excess allele sharing (Pearson’s r = 0.25-0.38) except for comparisons that tested excess allele sharing of different *mloto A* or *mloto C* populations with *mloto B*, which showed no significant correlation with geographic location (Fig. 4.b). Taken together, our results are consistent with the presence of three widely distributed groups of *C. mloto* that have (occasionally) been exchanging genetic material in places where they were in contact, but still retain their separate genetic identities.

### *C. mloto* populations show strong excess of rare genetic variants

Investigating population genetic summary statistics, we found relatively similar levels of nucleotide diversity among populations and clades (Fig. 5A, supplementary Table XI). The relatively largest variation is seen among *mloto C* populations with the relatively smallest value of 0.1336% ± 0.0139 for the Chilumba population and the largest value of 0.1414% ± 0.0149 for the Nankhwali trawler population. Unexpectedly, despite its peripheral geographic location and recent demographic changes due to fishing, the *mloto A* population from Lake Malombe (0.1398% ± 0.0141) showed one of the highest values of genetic diversity. With 0.1365% ± 0.0149, the estimated nucleotide diversity for *C. virginalis* was at the lower end of values measured for *C. mloto* populations.

**Figure 5.**
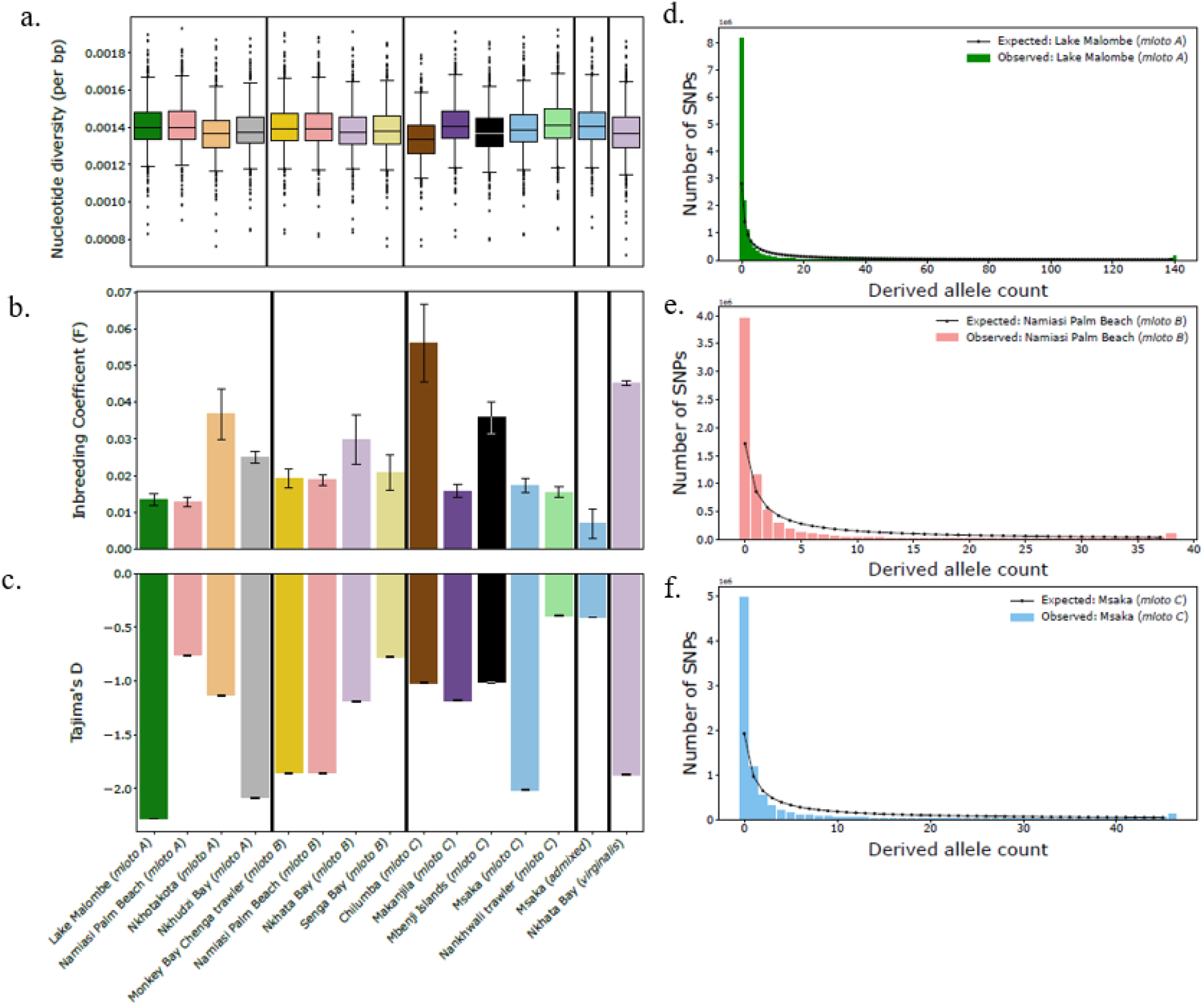
Population genetic summary statistics (a) Nucleotide diversity () per bp calculated in windows of 1Mbp. (b) Average inbreeding Coefficient (F) per population (with n>1). (c) Tajima’s D in windows of 100kb per population (with n>1). Observed allele frequency spectra for (d) the Lake Malombe population of clade *mloto A;* (E) the Namiasi palm beach population of clade *mloto B*, and (f) the Msaka population of clade *mloto C*. Black dots/lines show the respective expectations for neutrally evolving populations of constant size.

Inbreeding coefficients, measuring a deficiency of heterozygote genotypes relative to Hardy-Weinberg expectations, were moderately positive in all populations (Fig. 5B), consistent with either moderate degrees of (historic) mating between related individuals or population substructuring. However, the fact that relatively high inbreeding coefficients coincide with relatively low nucleotide diversity – the highest values most variable along the genome being seen in the least diverse Chilumba *mloto C* – suggests that inbreeding coefficients reflect within population demographic events rather than residual population structure. *C. virginalis* showed the second largest inbreeding coefficient with little variation of it along the genome, a signal which might point to a historic population bottleneck.

To gain further insight into how utaka genetic diversity patterns have been shaped by past demographic events, we computed Tajima’s D, a summary of the distribution of allele frequencies (Fig. 5C). For the population of each clade with largest sample size, we also computed full site frequency spectra (SFS) and compared them to expected spectra under neutral evolution (Fig. 5D-F). These analyses revealed an excess of rare genetic variants (negative Tajima’s D) compared to neutral expectations in all populations, with strongly negative Tajima’s D values close to -2 being found for populations across the *C. mloto* clades and also for *C. virginalis*. Such an excess of rare genetic variants can for example be caused by historic population expansion or by directional selection. That said, the presence of negative values of Tajima’s D at a genome-wide scale, without large genomic variation (small error bars in Fig. 5C), suggests that the observed strong excess of rare genetic variants is mainly driven by strong population expansion in the time frame of the (largely shared) coalescent history of present day populations, rather than by positive selection, which would lead to genomically more localised allele frequency changes.

### Colour of male breeding dress varies with genetic clade membership

To investigate potential morphological differences between the different clades, we qualitatively assessed variation in male breeding dress across *C. mloto* clades based on photographs (Fig. 6). Male breeding dress is a key trait in reproductive isolation of Malawi cichlids that often differs between sister species (Maan and Sefc, 2013). Unfortunately, we only had photos or preserved specimens available for a subset of the individuals, namely 17 males of clade *mloto A* (all except one from Lake Malombe), 20 males of clade *mloto C*, and only two males of *mloto B*. Although the present sample size is too small for quantitative conclusions, the data suggests that the general pattern of black body and yellow dorsal fin colouration of breeding males common to many utaka shows clade specific variation. Specifically, all examined males of *mloto A* showed clear yellow colouration in the dorsal fin covering the whole fin at the anterior end and extending relatively far towards the posterior end at the upper fin margin (while the lower margin has a blackish colour). At the same time, *mloto A males* generally featured a yellow blaze on the forehead above the eyes, a pattern that can be hard to spot on photographs but is very apparent in examined specimens. Therefore, we tentatively refer to the *mloto A* breeding dress morphotype as “Yellow Head Yellow Dorsal” (YHYD). Breeding males of *mloto C* showed a dorsal fin colouration generally similar to *mloto A*, albeit with more variation in the relative amounts of black and yellow, but they generally do not show any discernible yellow blaze on their forehead, a morphotype to which we tentatively refer as “Black Head Yellow Dorsal” (BHYD). Finally, the two *mloto B* in male breeding dress, one from Nankhwali and one from Chiweta, showed a strong yellow blaze on their forehead but no or only very little yellow in their dorsal fin, wherefore we refer to their morphotype as “Yellow Head Black Dorsal” (YHBD). In summary, we found tentative evidence for clade-specific variation in male breeding dress, but a more comprehensive set of matched sequence and phenotype data will be necessary to scrutinise morphological differentiation among genetic clades.

**Figure 6.**
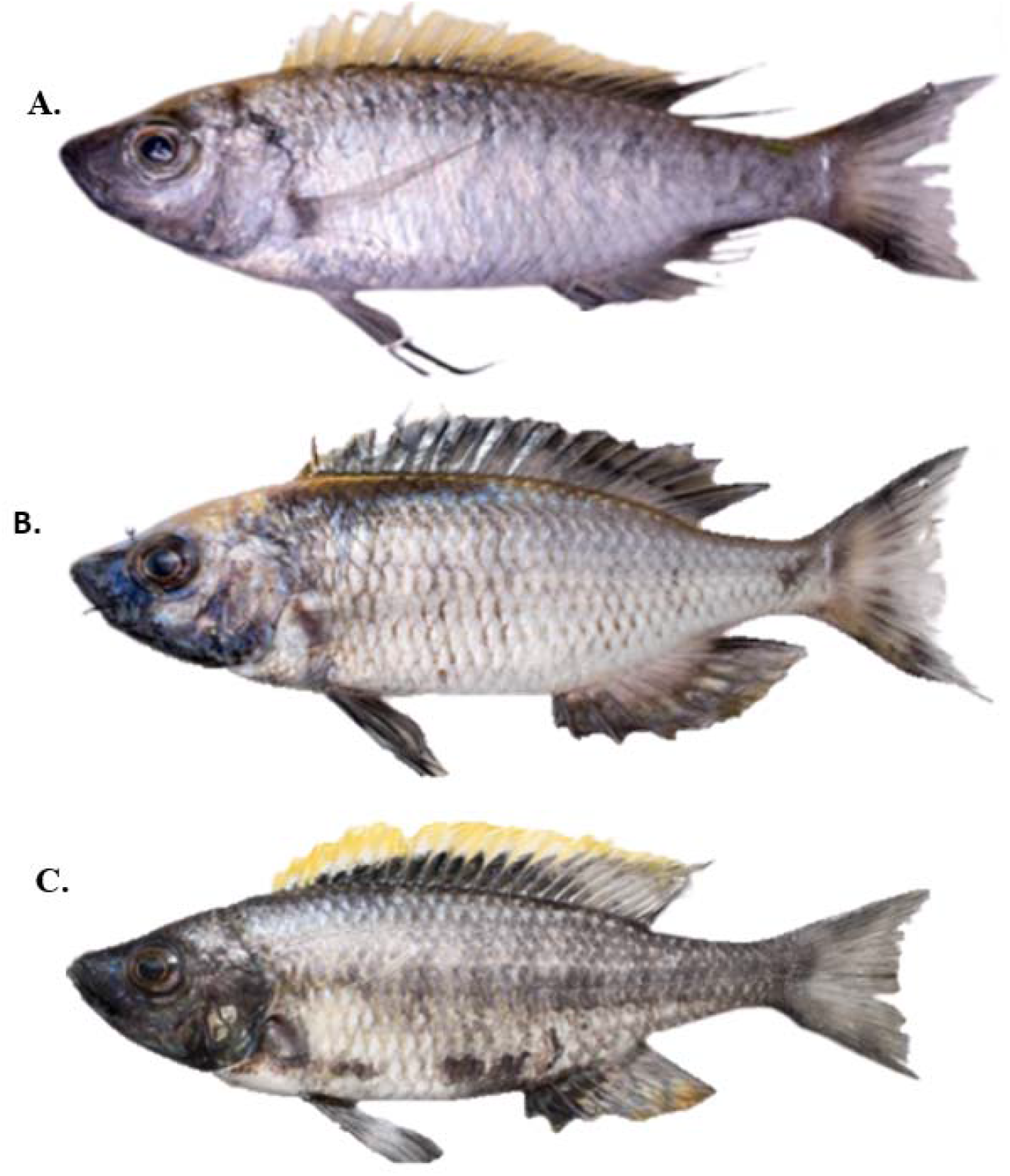
Representative males in breeding colouration in the three cryptic clades of *Copadichromis mloto*. A) *Copadichromis mloto* Yellow Head Yellow Dorsal (YHYD) “*mloto A* clade”, B) *Copadichromis mloto* Yellow Head Black Dorsal (YHBD) “*mloto B* clade”, C) *Copadichromis mloto* Black Head Yellow Dorsal (BHYD) “*mloto C* clade”.

## Discussion

An accurate understanding of genetic relationships, geographic distribution, and evolutionary history among populations plays a crucial role in the management of species of conservation concern. The pure utaka were recently assessed as near threatened due to a 40% decrease in catch rates between 1999 and 2016 likely due to intense fishing pressure (Konings, 2019). As we confirm here, this assessment probably mainly concerns *C. mloto* (previously also referred to as *C. sp*. ‘virginalis kajose’) – the pure utaka most common in artisanal and commercial catches and the main focus of this study. Here, we assess the population structure of *C. mloto* based on an almost lake-wide sample of whole genome sequences derived from fisheries catches and find – despite the general close relatedness typical for members of the Lake Malawi cichlid adaptive radiation – clear evidence for the presence of three genetically distinct groups. All three groups seem widely distributed with largely overlapping ranges and local populations that show evidence for variable levels of genetic exchange both with *C. virginalis* and with geographically proximate populations of other *C. mloto* groups. Together, this suggests that evolutionary relationships in the utaka are even more complicated than previously thought, with *C. mloto* alone deserving the recognition of at least three separate taxonomic entities.

The fact that, despite overlapping ranges, most specimens collected from a given fisheries catch pertained to the same *C. mloto* clade may indicate either a tendency to school according to species or that there is some kind of habitat partitioning, but the presence of exceptions to this trend and of apparently admixed individuals corroborates at least some degree of sympatry and occasional interbreeding. *C. mloto* are known to have defined breeding seasons when they aggregate in sandy or muddy habitats where males build and defend mating platforms (bowers), while outside of breeding season they follow a semi-pelagic lifestyle mainly feeding on plankton in inshore open-water habitats (Turner, 1996; Konings, 1999; Anseeuw et al. 2008). A difference in timing or location of breeding aggregations might contribute to the maintenance of differentiation between the clades, but further research will be needed to better understand ecology and life history of *C. mloto* groups – an endeavour timely and urgent given the worrisome conservation status of this species and continuing exploitation.

Despite the relatively isolated geographic position of Lake Malombe, being connected to Lake Malawi through the ∼15km long Upper Shire river, its *C. mloto* population clustered together with populations from Nkhudzi Bay in the south east arm and Nkhata Bay at the central east coast of Lake Malawi, together forming clade *mloto A*. Given that Lake Malombe had completely dried out and only refilled in the 1930s (Tweddle et al. 1995), it is possible that *C. mloto* from Lake Malombe are relatively recent migrants from Lake Malawi, which is consistent with observations that the Upper Shire River is used by different species to migrate between the lakes (FAO, 1993). However, the fact that, with the exception of a single specimen, the population from Namiasi Palm Beach collected less than four kilometres from Lake Malawi’s drainage into the Upper Shire belongs to a completely different genetic group (clade *mloto B*) suggest that migration through the Upper Shire is presently not a very common event or is specific to one clade. Also consistent with previous work (Anseeuw et al. 2011), we did not find any evidence for a population bottleneck in the founding of the Lake Malombe population – on the contrary, nucleotide diversity was relatively high and inbreeding coefficient at the low end of all populations. The strongly negative values of Tajima’s D in this population could reflect recent selection which would be consistent with the extreme fishing pressure reported for Lake Malombe (Chavula et al. 2023). However, further analysis will be necessary to disentangle evolutionary changes due to (fisheries-induced) selection from those brought about by demographic changes.

An important open question is whether the genetic differentiation of *C. mloto* clades is also reflected in morphological differences. Answering this question is complicated by the fact that only a fraction of the specimens used in this study have been preserved, that no 1:1 match of specimens and DNA samples is available for the 2002 collection, and that the large variation in adult specimen body size across locations (which might reflect variation in fishing pressure, see below) makes morphological examination tricky for the current sample set. That said, focusing on specimens in male breeding colouration that could be matched to DNA sequences we found some tentative evidence that variation in male breeding dress correlates with adherence to a genetic clade. This is worth further examination given that assortative mating based on male breeding colouration has been shown to be an important factor in the maintenance of species barriers in cichlids (Blais et al. 2009; Madeleine et al. 1998).

Previous analysis of the population structure of *C. mloto* based on microsatellites and mtDNA of a partly overlapping sample set found low but significant degrees of population substructuring of *C. mloto*, but was not able to identify more than a single genetic cluster (Anseeuw et al. 2008). Thus, while overall consistent, our results exemplify the increased power of whole genome data in identifying geographic substructuring among evolutionary recent lineages. While previous studies mainly reported population structure within species restricted to the rocky habitat (e.g., mbuna and *Protomelas* (Van Oppen at al., 1997; Arnegard et al. 1999; Pereyra et al. 2004), our results emphasise that sufficiently dense data, both in terms of genetic markers as well as geographic coverage, may reveal intricate patterns of population structure and previously unknown taxonomic diversity, also in sand dwelling and (semi) pelagic species.

Both decreasing catch rates and an observed decrease in the size of breeding adults suggest that fishing has had a profound effect on *C. mloto* (Anseeuw et al. 2011), affecting individual survival and potentially leading to fisheries-induced evolution (Heino et al. 2015). Here, we found that estimates of genetic parameters do not yet reflect demographic and life history changes brought about by fishing: *C. mloto* shows relatively higher diversity and generally lower inbreeding coefficients compared to the probably less fished *C. virginalis*, and relatively little geographic variation in genetic diversity despite strong observed variation in local fishing pressure. On the contrary, the remarkably strong bias towards rare genetic variants in all examined *C. mloto* and *C. virginalis* populations (and consequently strongly negative Tajima’s D) points to a strong historic population expansion. This signal might reflect a rapid expansion of Utaka into open water habitats at the time such habitats became available after the lake recovered from almost complete desiccation ∼75 kya (Ivory et al. 2016; Salzburger et al. 2014). This is consistent with studies suggesting that wet-dry transitions in Lake Malawi over the past 1.2 million years and resulting habitat variability have shaped the rapid diversification of Malawi cichlids through changing niche availability and colonisation (Ivory et al. 2016). Combining detailed demographic modelling with paleobiological data (e.g., on lake level fluctuations) will yield a more precise understanding of utaka’s evolutionary history and diversification (Danley et al. 2000; Sturmbauer et al. 2001; Nevado et al. 2013; Ivory et al. 2016).

The lack of population genetic signal for fisheries-induced demographic changes is not unexpected given that the onset of fishing pressure is relatively recent on evolutionary scales and that the mentioned population statistics summarise demographic patterns over long timescales. Indeed, theoretical research suggests that although fishing pressure strongly reduces the current effective population size, the effects of 100 years of intense fishing on genetic diversity are minor (Marty et al. 2015). Earlier studies of fisheries-induced evolution in marine systems have not been able to find genetic evidence for fisheries-induced evolution, attributing observed life history changes to phenotypic plasticity (Jørgensen et al. 2009; Enberg et al. 2012). We suggest that larger sample sizes and bespoke statistics will be necessary to assess the within-population effect of fishing on patterns of genetic diversity. That said, spatially variable fishing pressures are also expected to engender changes in the relative abundance of different species and genetic clades, a signal that might be easier to detect than within-population demographic and evolutionary effects. Specifically, fishing could affect the relative abundance of *C. mloto* populations pertaining to different genetic clades. Sufficiently fine-scaled geographic and temporal genetic monitoring of *C. mloto* populations and other commercially important Malawi cichlid species more generally will be crucial for the timely detection of such changes and appropriate management responses.

## Conclusion

We investigated population genetic structure in the Malawi cichlid species *C. mloto* based on whole-genome data of a relatively large and geographically widespread sample set. We found that what was previously thought to be a single species of utaka – in itself a complex of semi-open water, planktivorous cichlid species – shows unexpected patterns of population structure, pointing to the presence of at least three widely distributed, genetically distinct *C. mloto* clades. Generalising from this, our results indicate that taxonomic diversity in the large and understudied group of benthic and pelagic Lake Malawi cichlid species might be higher and more complex than previously thought, despite generally wide distribution ranges and seemingly continuous habitats. Such diversity might often not be easily discernible by morphological examination and calls for broad genomic investigation and monitoring. This is especially urgent in the context of increasing commercial exploitation of most benthic and pelagic species.

## Acknowledgements

We would like to thank the Department of Fisheries in Malawi for permission to conduct fieldwork for the 2016 and 2017 sample collections and Mexford Mulumpwa, Milan Malinsky, Alix Tyers, Mingliu Du, Gregoire Vernaz, Karl Svardal, and Richard Zatha for their assistance during fieldwork. We sincerely thank Eric Miska and Richard Durbin (University of Cambridge and Wellcome Sanger Institute) for funding sample collection and sequencing for this study. We thank Jos Snoeks (Royal Belgian Museum for Central Africa), Erik Verheyen (Royal Belgian Institute of Natural Sciences) and Dieter Anseeuw for sharing fin clips and extracts of the 2002 *C. mloto* collection, and Maarten Van Steenbergen for help with locating samples.

## Authors contributions

W.S. and H.S. conceived the study. A.H. generated and prepared variant call format (VCF) data. W.S. analysed genomic data with contributions from A.H and H.S. M.G., B.R. and H.S.2 facilitated sample collection. S.G. processed sample metadata and performed DNA extractions. G.F.T undertook comparison of male breeding dress against clade membership. W.S. and H.S. wrote the manuscript with input from A.H. and G.F.T. All authors read and approved the manuscript.

## Data availability

Supporting data is made available in the online supplementary on an open-access basis for research use only. ENA Accession numbers of raw sequencing data will be added before final publication. Data was collected under appropriate ethical and sampling permits and genetic material and sequences are subject to an Access and Benefits Sharing (ABS) agreement with the Government of Malawi. Any person who wishes to use this data for any form of commercial purpose must first enter into a commercial licensing and benefit-sharing arrangement with the Government of Malawi.

