## Supplementary material 1 for "Population genomic analysis reveals cryptic population structure in the commercially important Lake Malawi cichlid *Copadichromis mloto*"

**Supplementary Table II.** Excluded chromosome inversion regions in the genome sequences of *Copadichromis mloto*/*C. virginalis*

| Chromosome | Excluded inversions regions (bp) |
| --- | --- |
| Chr 2 | 9,325,000 – 33,010,000 |
| Chr 9 | 11,500,000 – 29,400,000 |
| Chr 10 | 10,825,000 – 29,785,000 |
| Chr 11 | 6,270,000 – 29,210,000 |
| Chr 13 | 9,310,000 – 30,070,000 |

**Supplementary Fig. III*.*** Population admixture/structure analysis in *Copadichromis mloto/virginalis* shows three main strong separate colour proportions distribution relationship of individuals in *C. mloto and C. virginalis*.

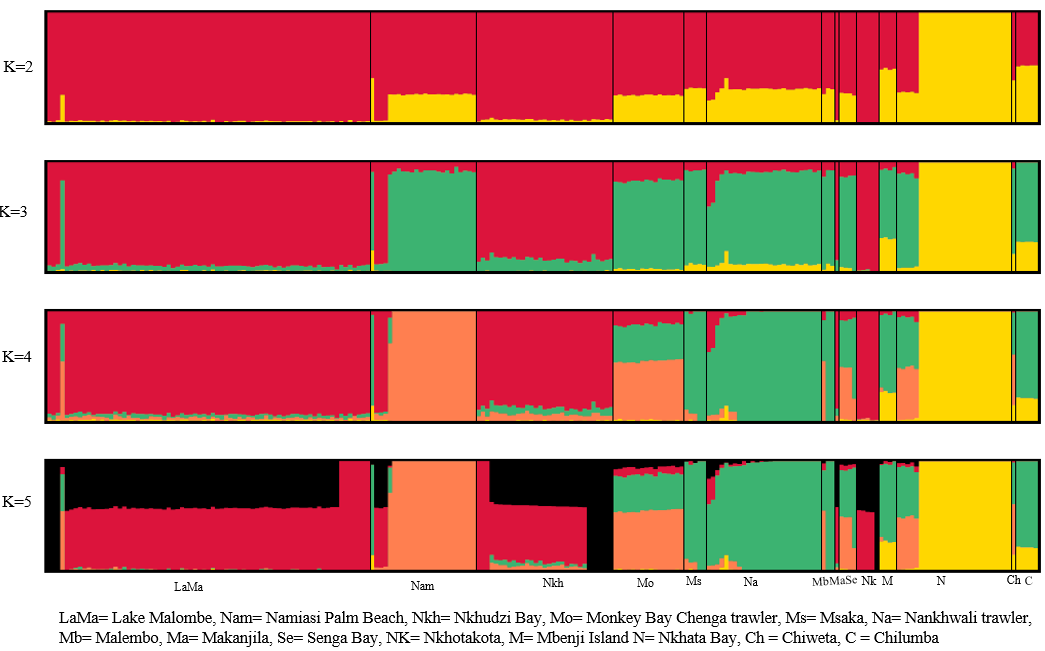

**Supplementary Fig. IV*.*** Population admixture/structure analysis in *Copadichromis mloto*

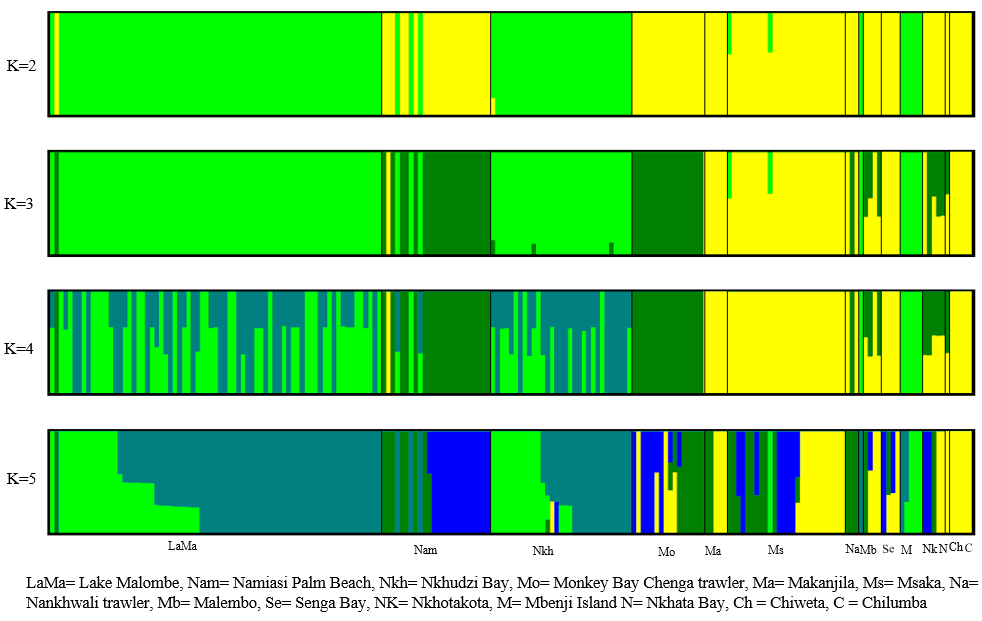

**Supplementary Fig. V.** Principal component analysis in *Copadichromis mloto* and *C. virginalis* showing the relationship among the species.

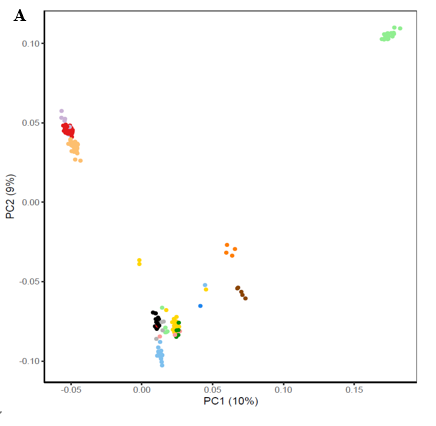

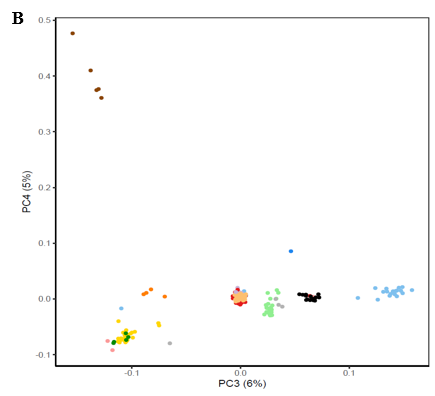

**Supplementary Fig. VI*.*** Scatter plot showing the number of missing genotypes (y-axis) in *Copadichromis mloto* and *C. virginalis* individuals (y-axis) plotted against depth of coverage (x-axis).

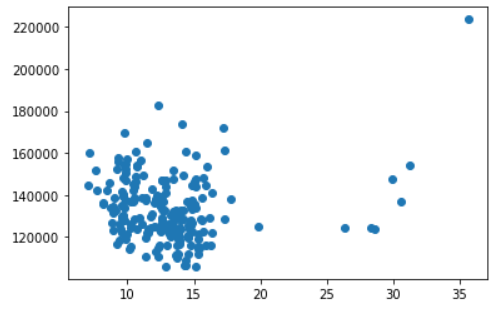

**Supplementary Fig. VII*.*** Scatter plot showing the number of singletons in *Copadichromis mloto* and *C. virginalis* individuals (y-axis) plotted against depth of coverage (x-axis).

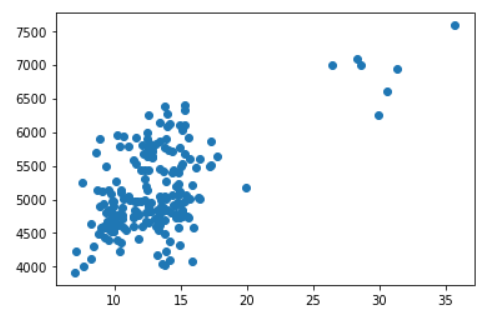

**Supplementary Fig. VIII*.*** Number of missing genotypes and number of reference homozygosity runs (nReFHom) in *Copadichromis mloto* and *C. virginalis* individuals

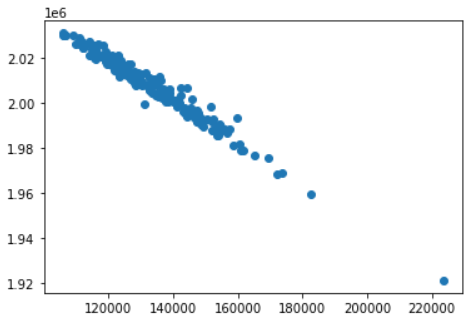

**Supplementary Table IX.** Cross-validation error values for admixture analysis of *Copadichromis virginalis and C. mloto* with a range of K-values from 2 to 5.

| K-value | Cross-validation error |
| --- | --- |
| 2 | 0.46833 |
| 3 | 0.46805 |
| 4 | 0.47589 |
| 5 | 0.48948 |

Cross-validation plot for Copadchromis virginalis & C. mloto

CV error values

| K-values | Cross-validation error |
| --- | --- |
| 2 | 0.46833 |
| 3 | 0.46805 |
| 4 | 0.47589 |
| 5 | 0.48948 |

Cross-validation plot of admixture K-values in the range from 2 to 5 of *Copadichromis virginalis* and *C. mloto*

**Supplementary Table X.** Cross-validation error values for admixture analysis of *Copadichromis mloto* with a range of K-values from 2 to 5.

| K-value | Cross-validation error |
| --- | --- |
| 2 | 0.22628 |
| 3 | 0.23962 |
| 4 | 0.25344 |
| 5 | 0.26691 |

Cross-validation plot for Copadichromis mloto

CV error values

| K- vlalues | Cross-validation error |
| --- | --- |
| 2 | 0.22628 |
| 3 | 0.23962 |
| 4 | 0.25344 |
| 5 | 0.26691 |

Cross-validation plot of admixture K-values in the range from 2 to 5 of *Copadichromis mloto*

**Supplementary Table XI.** Nucleotide diversity per site measurements in the three cryptic clades (*mloto A/B/C*), admixed populations and *C. virginalis* represented in median and standard error values expressed in percentage

| Name of the clade | Nucleotide diversity (median and standard deviation values) expressed in percentage |
| --- | --- |
| Lake Malombe *mloto A* | 0.1398 ± 0.0141 |
| Namiasi Palm Beach *mloto A* | 0.1334 ± 0.0141 |
| Nkhotakota *mloto A* | 0.1398 ± 0.0140 |
| Nkhudzi Bay *mloto A* | 0.1373 ± 0.0139 |
| Monkey Bay Chenga trawler *mloto A* | 0.1395 ± 0.0143 |
| Namiasi Palm Beach *mloto B* | 0.1394 ± 0.0144 |
| Nkhata Bay *mloto B* | 0.1375 ± 0.0142 |
| Senga Bay *mloto B* | 0.1379 ± 0.0144 |
| Chilumba *mloto C* | 0.1336 ± 0.0139 |
| Makanjila *mloto C* | 0.1407 ± 0.0145 |
| Mbenji Islands *mloto C* | 0.1369 ± 0.0139 |
| Msaka *mloto C* | 0.1386 ± 0.0143 |
| Nankhwali trawler *mloto C* | 0.1414 ± 0.0149 |
| Msaka admixed | 0.1402 ± 0.0145 |
| Nkhata Bay virginalis | 0.1365 ± 0.0149 |
