## Supplementary figures and images for "Population genomic analysis reveals cryptic population structure in the commercially important Lake Malawi cichlid *Copadichromis mloto*"

### Supplementary material 2

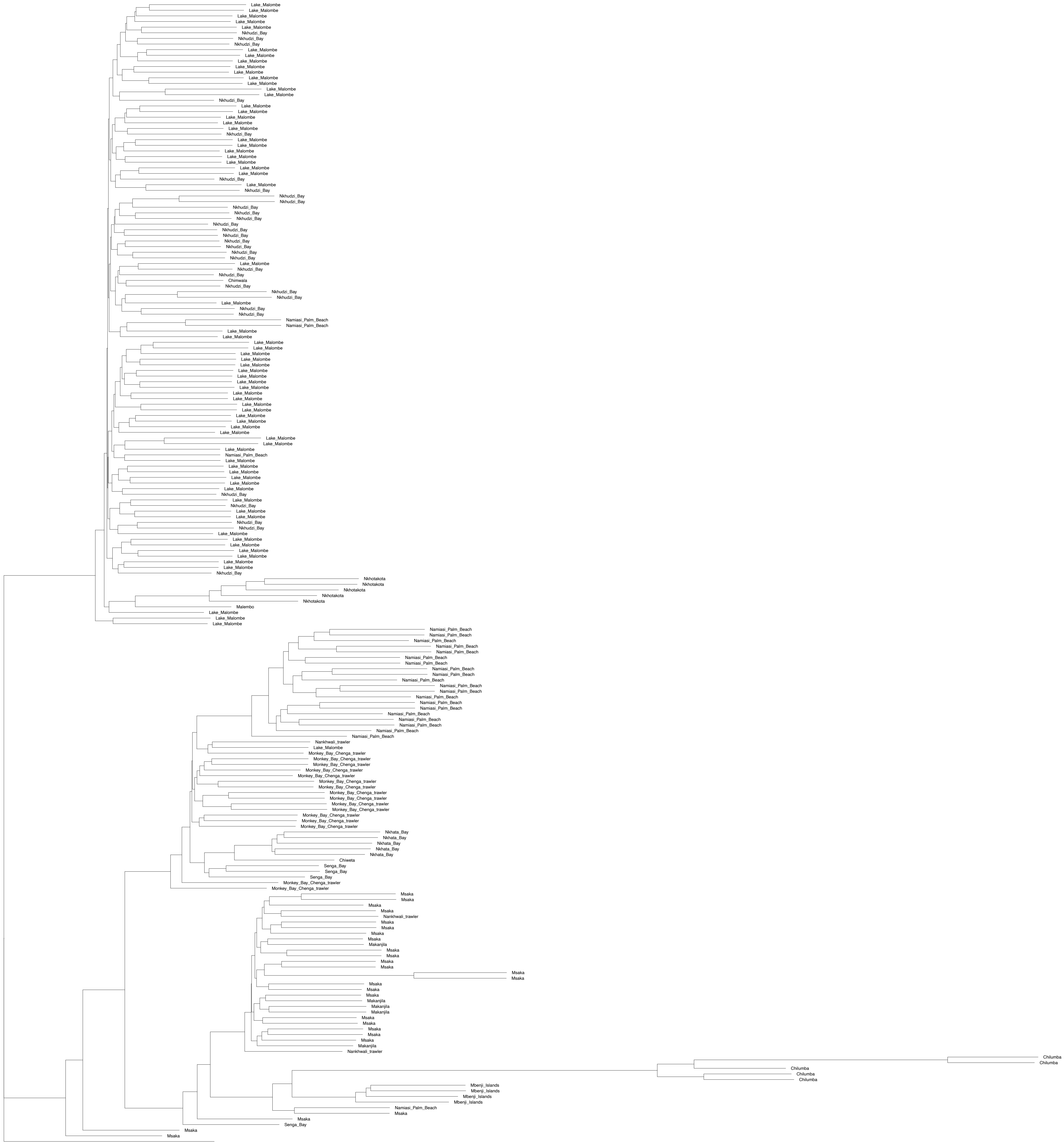
